# The CATION CALCIUM EXCHANGER 4 (CCX4) regulates LRX1-related root hair development through Ca^2+^ homeostasis

**DOI:** 10.1101/2025.06.25.660713

**Authors:** Xiaoyu Hou, Giorgia Tortora, Aline Herger, Stefano Buratti, Petre Dobrev, Roberta Vaculiková, Jozef Lacek, Alexandros Georgios Sotiropoulos, Gabor Kadler, Myriam Schaufelberger, Alessia Candeo, Andrea Bassi, Thomas Wicker, Alex Costa, Christoph Ringli

## Abstract

Calcium, as a cellular second messenger, is essential for plant growth. A tip-focused Ca^2+^ gradient in polarized cells is considered to drive cell expansion. The cell wall polysaccharide pectin is a major Ca^2+^ binding structure and Ca^2+^ homeostasis is influenced by the cell wall architecture. LRR-extensin (LRX) proteins are extracellular regulators of cell wall development that are anchored in the cell wall by their extensin domain. The extensin-less *LRX1ΔE14* variant of the root hair-expressed *LRX1* of Arabidopsis induces a dominant-negative effect resulting in aberrant root hairs. In an effort to identify the underlying mechanism of the root hair defect caused by *LRX1ΔE14*, we isolated a *<u>su</u>ppressor of dominant-<u>ne</u>gative effect* mutant, *sune42*. It codes for the CATION CALCIUM EXCHANGER 4 (CCX4) that localizes to the Golgi apparatus and was shown to have Ca^2+^ transport activity. A detailed investigation of the Ca^2+^ dynamics revealed that LRX1ΔE14 coincides with a defect in tip-focused cytoplasmic Ca^2+^ oscillation, and this effect is alleviated by the *sune42* mutation. Additionally, reducing Ca^2+^ availability influences the *LRX1ΔE14*-induced root hair defect. We conclude that *sune42* suppresses the root hair defect in *LRX1ΔE14* through modulating cytoplasmic Ca^2+^ dynamics, pointing at the importance of the Golgi apparatus for cellular Ca^2+^ homeostasis.

## Introduction

Plant cell growth is constrained by the cell wall, which undergoes periodic structural modifications to allow the cell to expand. The cell wall is a complex network of cellulose microfibrils embedded within a matrix of polysaccharides, interwoven with proteinaceous molecules (Cosgrove, 2024). The cell wall-localized Leucine-rich repeat extensins (LRXs) are chimeric proteins that contain a C-terminal extensin domain and an N-terminal LRR domain, separated by a cysteine-rich domain (Rubinstein et al., 1995; Baumberger et al., 2001; Ringli, 2010; Herger et al., 2019). LRXs influence cell wall formation by binding RALF (rapid alkalinization factor) peptides and compacting pectin (Mecchia et al., 2017; Moussu et al., 2020; Moussu et al., 2023; Schoenaers et al., 2024). Genome analysis revealed the presence of eleven *LRXs* in *Arabidopsis*, among which *LRX1* and *LRX2* are predominantly expressed in root hairs (Baumberger et al., 2003a). Loss of *LRX1* function results in shorter root hairs that often branch or burst (Baumberger et al., 2001; Baumberger et al., 2003). The extensin domain of LRXs, which contains Ser-Hyp_n_ repetitive sequences characteristic for structural hydroxyproline-rich glycoproteins (HRGPs) (Borassi et al., 2016; Liu et al., 2016), anchors the protein in the extracellular matrix. It is essential for the function of LRX1, as LRX1 lacking the extensin domain (LRX1ΔE14) is unable to complement the *lrx1* root hair phenotype (Ringli, 2010). Furthermore, the expression of extensin-less *LRXs* in wild-type Col plants induces a dominant-negative effect (Ringli, 2010; Dünser et al., 2019).

Refined integration of extracellular and cytoplasmic events is needed for coordinating between cell wall enlargement and cell expansion. LRXs have been suggested to function in conjunction with the *Catharanthus roseus* Receptor-Like Kinase1-Like protein (*Cr*RLK1L) FERONIA (FER) to translate the extracellular dynamics into cytoplasmic responses. *lrx* mutants phenocopy the *fer-4* mutant, which shows root hair impairment similar to the *lrx1 lrx2* mutant (Herger et al., 2020) and shoot growth defects similar to the *lrx3 lrx4 lrx5* mutant (Draeger et al., 2015; Zhao et al., 2018; Dünser et al., 2019). The multitude of processes influenced by the LRX-RALF-FER module is only beginning to be understood (Leiber et al., 2010; Schaufelberger et al., 2019; Song et al., 2022; Wang et al., 2022; Pacheco et al., 2023; Gupta et al., 2024).

Tip-growing cells, like pollen tubes and root hairs, are featured for their rapid and unidirectional growth. Polarized cell growth relies on the apical fusion of secretory vesicles, which is under the control of the configuration of the actin cytoskeleton (Campanoni and Blatt, 2007; Ketelaar, 2013). This process is intimately linked with Ca^2+^ gradients and Ca^2+^ oscillations in the cytoplasm (Wymer et al., 1997; Zhang et al., 2017). Mutations in *FER* and several other *CrRLK1L* family members such as *ERULUS* and *ANX1/2* dampen the tip-focused cytoplasmic Ca^2+^ gradients in tip-growing cells (Ngo et al., 2014; Kwon et al., 2018; Schoenaers et al., 2018; Gao et al., 2023; Schoenaers et al., 2024). A few Ca^2+^ channels involved in CrRLK1L-mediated Ca^2+^ signaling, like powdery mildew resistance locus (MLOs), have also been identified (Ogawa et al., 2025). Other studies have revealed the essential roles of several cyclic nucleotide-gated channels (CNGCs) in polarized growth by modulating tip-focused Ca^2+^ oscillations, among which CNGC6, CNGC9, and CNGC14 act together in controlling root hair polarity and integrity (Zhang et al., 2017; Brost et al., 2019; Zhu et al., 2025).

Importantly, organelles have been found to largely contribute to the cytoplasmic Ca^2+^ signatures (Trewavas et al., 1996; Kudla et al., 2010; Costa et al., 2018; Resentini et al., 2021). Organellar channels, transporters, and pumps involved in Ca^2+^ transport (e.g. cation/Ca^2+^ exchangers (CCXs), Ca^2+^-ATPases) have been reported to regulate a range of physiological processes (Bose et al., 2011; Frei dit Frey et al., 2012; Pittman and Hirschi, 2016; Corso et al., 2018; Costa et al., 2023; Kanamori et al., 2023). In plants, the Golgi apparatus is known to be essential for protein posttranslational modification and trafficking (Vitale and Galili, 2001) and cell wall polysaccharide synthesis (Cosgrove, 2024; Gupta et al., 2024). These processes are tightly linked with Ca^2+^, but little is known about Ca^2+^ homeostasis and signaling in the Golgi apparatus (Costa et al., 2018; Pirayesh et al., 2021). The estimated resting free Ca^2+^ in the Golgi is slightly higher than that in the cytoplasm and transient Ca^2+^ changes in the Golgi during stress responses have been observed (Ordenes et al., 2012). A recent study has shown that a Golgi-localized CCX member in *Arabidopsis*, CCX4, mediates Ca^2+^ transport and is involved in Ca^2+^ response and osmotic tolerance (Kanamori et al., 2023).

To seek for novel modulators of the LRX1-related signaling pathway, a suppressor screen was performed on the *LRX1ΔE14*-expressing line that displays a defect in root hair development. This led to the identification of *sune42* (*suppressor of dominant-negative effect of LRX1ΔE14_42*), which harbors a mutation in the gene *Cation Calcium Exchanger 4* (*CCX4*). We found that the previously reported role of *CCX4* in influencing NPR1-dependent salicylic acid (SA) signaling (Fujikura et al., 2020) is not involved in suppressing the root hair defect in *LRX1ΔE14.* Our study shows that *LRX1ΔE14* is impaired in Ca^2+^ oscillation and mutation in *CCX4* likely alleviates this derangement induced by *LRX1ΔE14*. Our data imply that *sune42* mutation might restore the *LRX1ΔE14* root hair phenotype through regulating Ca^2+^ dynamics by altering the flux of Ca^2+^ between the Golgi and the cytoplasm.

## Materials and Methods

### Plant material and growth conditions

*Arabidopsis thaliana* Col was used for all experiments. The p*LRX1*::*LRX1ΔE14* line (referred to *LRX1ΔE14*) is described in Ringli, 2010. *sune42* (in Col), *lrx1 sune42*, *fer-5 sune42* were obtained by crossing *sune42* with wild-type Col, *lrx1* (Baumberger et al., 2001), and *fer-5* (Duan et al., 2010), respectively. *ccx4* and *npr1* were generated by *Crispr CAS9*-guided mutagenesis. *LRX1ΔE14 sune42* was later crossed with *npr1* to produce *LRX1ΔE14 sune42 npr1.* For the cytoplasmic calcium measurement, the transgenic line containing *pUBQ10::R-GECO1* transgene (Keinath et al., 2015) was crossed with *sune42*, *LRX1ΔE14, LRX1ΔE14 sune42*, and the resulting F2 populations were screened by phenotype and molecular markers for the individual mutations and transgenes.

Unless otherwise detailed, *Arabidopsis* seeds were surface sterilized with 1% (v/v) sodium chlorite, 0.03% (v/v) Triton X-100 and washed three times with sterile water. After plating, they were stratified for 2 days at 4°C in darkness and subsequently grown vertically on ½ MS/Vitamins with 2% (w/v) sucrose, 10 mg/L myo-inositol, 0.5 g/L MES pH 5.7, and 0.6% (w/v) Gelrite (Duchefa). All seedlings were grown in a long-day regime (16-h light/8-h dark) at 22°C. Root hairs were visualized 5 days after germination. For propagation, transformation, and crossing, seedlings were transferred to soil and grown in the growth chamber at 22°C with a 16-h light/8-h dark cycle.

### Plasmid construction

For *CRISPR Cas9* mutagenesis, the *pKI1.1* (Tsutsui and Higashiyama, 2016) plasmid digested with *AarI* was ligated with the double-stranded oligos CCX4_gRNA_F/R targeting the *CCX4* gene and NPR1_gRNA_F/R targeting *NPR1* (Table 1). For the complementation assay, the *CCX4* promoter and coding sequence of *CCX4* were separately amplified using the primers SpeI_CCX4_F and CCX4_BamHI_R for fragment 1, primers CCX4_BamHI_F and CCX4_AscI_R for fragment 2 (Table 1), and cloned into pJET vector with *SpeI/BamHI* and *BamHI/AscI*, respectively. After sequencing, correct clones of the two fragments were fused by a *BamHI* restriction site in both clones to generate *pJET-pCCX4::CCX4*. *pCCX4::CCX4* was then cut from *pJET* with *SpeI* and *AscI* and cloned into *pGPTV-Bar-ocsTerminator* (Becker et al., 1992).

**Table 1.**
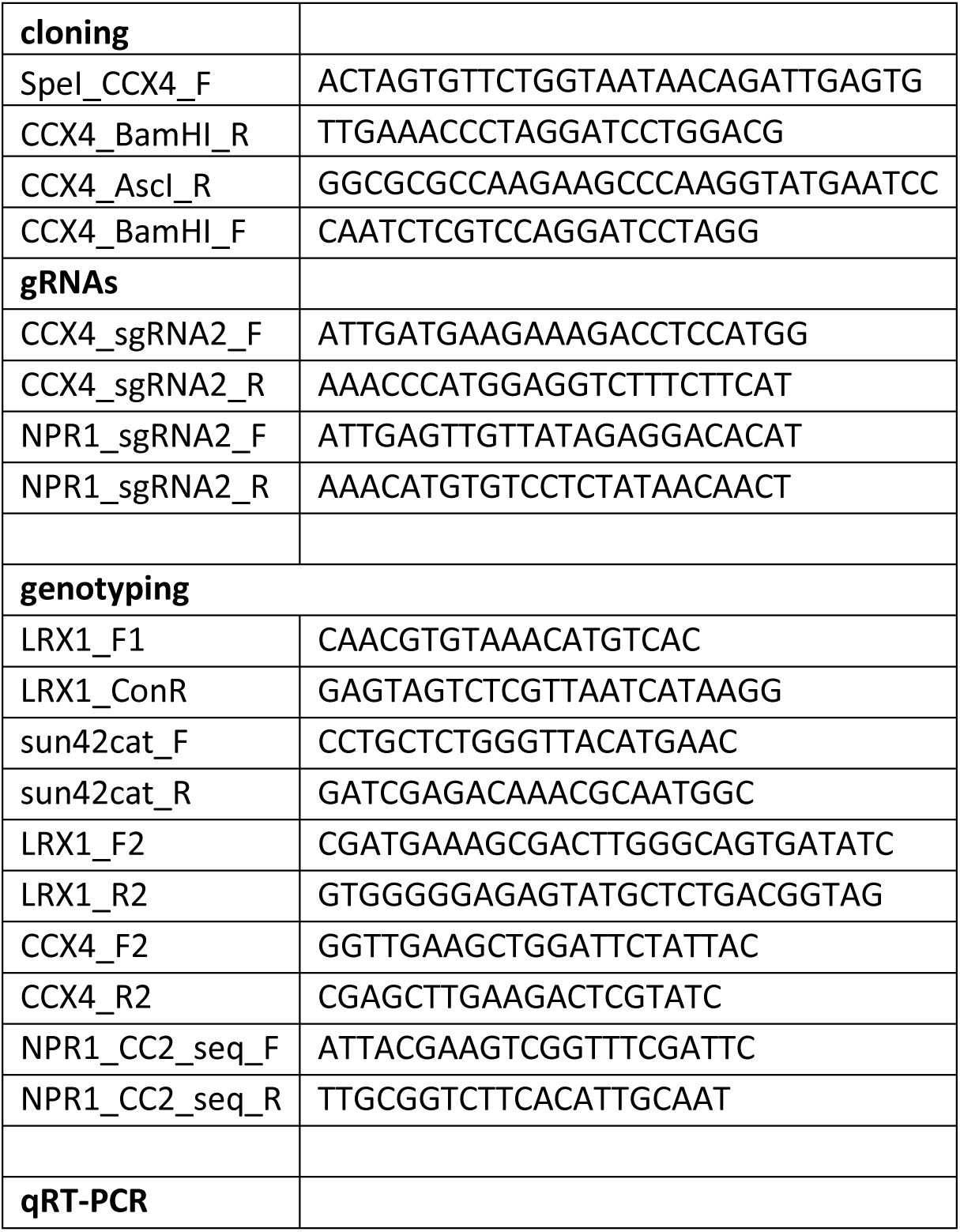
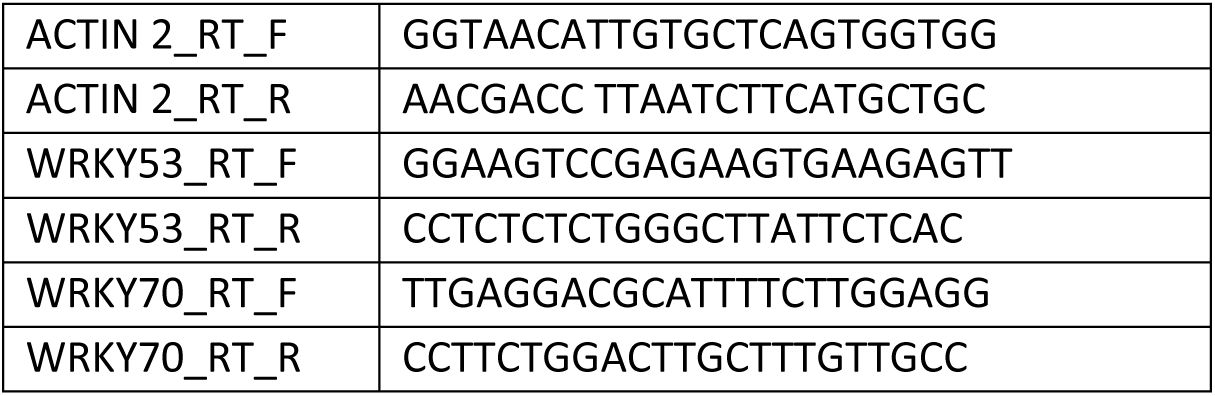
Primers used in this study.

### EMS mutagenesis

The ethyl methanesulfonate (EMS) mutagenesis and whole genomic sequencing was performed as described elsewhere (Guérin et al., 2025). Seeds from a line expressing *LRX1::LRX1ΔE14* were incubated in 100 mM phosphate buffer overnight. The next day, seeds were incubated in 100 mM phosphate buffer containing 0.2% EMS for 8 h on a shaker. The M1 seeds were rinsed 15 times with 300 mL of water and then grown directly on soil in 240 pots, each containing 10 plants, to propagate to the M2 generation. Each pot represents a batch of M2 seeds. On average, around 20 seeds per M2 plant (200 seeds per batch) were screened for seedlings with a suppressed *LRX1ΔE14* root hair phenotype. Selected putative mutant M2 seedlings were propagated and confirmed in the M3 generation. Positive lines were crossed with the non-mutagenized parental *LRX1ΔE14* line and propagated to the F2 generation that was analysed for segregation of the *sune* mutant phenotype.

### Whole genome sequencing, CAPS marker design

Whole-genome sequencing was performed as described in (Guérin et al., 2025). Ten F2 seedlings from the first backcross of *LRX1ΔE14 sune42* with the parental *LRX1ΔE14* line exhibiting a *LRX1ΔE14 sune42* phenotype were selected and pooled for DNA extraction. Whole-genome sequencing of the DNA of *LRX1ΔE14 sune42*, along with the non-mutagenized *LRX1ΔE14* line, was performed at Novogene using Illumina short-read technology. Raw sequence reads from the pooled *LRX1ΔE14 sune42* mutants were trimmed with the Trimmomatic (version 0.38) with the parameters LEADING:10, TRAILING:10 SLIDINGWINDOW:5:10, MINLEN:50. Trimmed sequence reads were mapped with the bwa software (version 0.7.17-r118) to the *Arabidopsis* Columbia reference genome (TAIR version 10) using default parameters. Resulting BAM files were sorted and duplicates removed with samtools (version 1.9). New read groups were assigned to the reads with the Picard software (version:2.27.5). Sequence variants were called with the GATK software (version 4.2). The vcftools software (version 0.1.16) was used to filter the vcf files using the parameters (--max-meanDP 7 –remove-indels). The analysis revealed three SNPs (single nucleotide polymorphisms) on chromosome 1 linked with the *sune42* mutation. Using this information, CAPS (cleaved amplified polymorphic sequences) markers were established, and co-segregation of the SNPs with the *LRX1ΔE14 sune42* phenotype was analysed.

For sequencing of the *LRX1ΔE14* construct in the identified *sune* mutants, the construct was PCR-amplified using *LRX1_F1* and *LRX1_TermR* primers (Table 1), targeting the promoter and terminator of LRX1, respectively. Due to the repetitive nature and length of the extensin coding sequence (Herger et al., 2019), the endogenous *LRX1* was not amplified.

For selection of *sune42* plants lacking *LRX1ΔE14*, a segregating F2 population of a backcross of *LRX1ΔE14 sune42* x Col was selected by PCR with the primers *LRX1_F1* and *LRX1_TermR* that detect *LRX1ΔE14* (Table 1).

### Plant transformation and selection

*Arabidopsis* transgenic lines were obtained by standard floral dipping method mediated by *Agrobacterium tumefaciens* (strain GV3101). For the complementation lines, T1 seeds of plants transformed with *pCCX4::CCX4* in *pGPTV-Bar* were selected on Basta selection media. The *CRISPR/Cas9* vector *pKI1.1R* (Tsutsui and Higashiyama, 2017) carries a *FASTRED* seed selection marker allowing selection based on RFP fluorescence. DNA from the cauline leaves of the inflorescences of T1 plants were extracted for genotyping using the primers CCX4_F2/R2 for *ccx4* (Table 1) and primers NPR1_F/R for *npr1*.

To detect the *sune42* mutation, the PCR product of *sune42cat_F* and *sune42cat_R* was digested with SacI, which only cuts the wild type. The *ccx4^crispr^* allele was detected using the primers *CCX4_F2* and *CCX4_R2*, and the product of only the wild type is cut by *NcoI*. The *lrx1* mutation was detected by PCR using the primers *LRX1_F2* and *LRX1_R2*, followed by a digestion with *EagI*, which only cuts the mutant. The *npr1^crispr^* allele was detected by sequencing the PCR product obtained with the primers *NPR1_CC2_seq_F* and *NPR1_CC2_seq_R*.

### Liquid chromatography-electrospray ionization tandem mass spectrometry (LC-ESI-MS/MS)

Phytohormone analysis was performed as described by Schmidt et al. (2024). Briefly, approximately 20 mg Fresh Weight of plant tissue was homogenized with 1.5 mm zirconium oxide beads using a FastPrep-24 instrument (MP Biomedicals) with 100 µL of 1 M HCOOH in water and a mixture of internal standards. After centrifugation at 17500 rpm for 20 minutes and re-extraction with additional 100 µL of 1 M HCOOH, the combined supernatant was applied to an SPE Oasis HLB 10 mg 96-well plate (Waters) pre-equilibrated with 100 µL of acetonitrile followed by the same volume of water and 1 M HCOOH. The supernatant was pushed through the SPE plate using a Pressure+96 manifold (Biotage). The 96-well SPE plate was washed three times with 100 µl of water. Samples were eluted with 100 µL 50% of acetonitrile/water (v/v). An aliquot of the eluate was injected into the LCMS system consisting of a UHPLC 1290 Infinity II (Agilent) coupled to a 6495 Triple Quadrupole mass spectrometer (Agilent). MS analysis was performed in MRM mode, two transitions per compound, using an isotope dilution method. Data acquisition and processing were performed using Mass Hunter software B.08 (Agilent).

### RNA extraction and qRT-PCR

mRNA extraction was done with the dynabeads mRNA DIRECT Kit (Invitrogen) according to the manufacturer’s protocol, with 25 µL of oligo(dT)25 per extraction. Roots of 10-day-old seedlings were frozen and ground in liquid nitrogen immediately upon collection. Around 20 mg of root tissue was used for each sample and 4 biological replicates of each treatment were included.

SA-responsive gene expression levels were assessed by qRT-PCR, with a CFX96 Real-Time System C1000TM Thermal cycler (Bio-Rad). Around 40 ng of mRNA was reverse transcribed using the iScript advanced kit (BioRad). qRT-PCR primers target *WRKY53* (WRKY53_F/R) and *WRKY70* (WRKY70_F/R) and the reference gene *ACTIN2* (ACTIN_F/R) are listed in Table 1. Four μL of 20-fold diluted cDNA was used for a total volume of 10 μL qRT-PCR reaction using KAPA SYBR FAST qPCR Master Mix (KK4601, Sigma–Aldrich) and 250 µM of each primer. Three technical replicates per sample were included.

### Root growth assay

Surface-sterilized seeds were grown on ½ MS agar plates supplemented with 2% sucrose, containing the chemical or corresponding solvent as control. For ethylene glycol-bis(β-aminoethyl ether)-N,N,N′,N′-tetra-acetic acid (EGTA) treatment, 1 mM EGTA was used, whereas for the salicylic acid (SA) treatment, 10 μM SA in EtOH was used and EtOH was added to the media of the control group. Root hair phenotypes were visualized after 5 days of growth. The plates were scanned and root (hair) length was measured by Fiji imageJ (Schindelin et al., 2012).

### Microscopy and quantitative imaging analysis

Experiments were conducted over a period of two weeks using two different sets of samples prepared according to the protocol described in (Romano Armada et al., 2019). The fluorescence light sheet setup implemented is a modified version of the single-sided illumination setup described in (Candeo et al., 2017). In the current setup, a laser at 561 nm is used as excitation source for the creation of the light sheet illumination. The detection unit, held orthogonally to the excitation axis, has been optimised to perform imaging of samples encoding the intensiometric calcium indicator R-GECO1 and includes a 10X water-dipping microscope objective (NA=0.3, UMPLFLN 10X W, Olympus) directly followed by a long pass filter at 600 nm (FELH0600 - Ø25.0 mm Longpass Filter, Cut-On Wavelength: 600 nm, Thorlabs), a tube lens (U-TLU-1-2, Olympus) and the camera (Neo 5.5 sCMOS, 2560×2160 pixels, ANDOR). Experiments were performed according to the protocol described in (Candeo et al., 2017). For each sample, 270 time points were recorded with a sampling time of 3 seconds acquiring 15 planes spaced 3 µm apart for each time point with an exposure time of 100 ms. Maximum Intensity Projection (MIP) of the 15 slices was then computed for each time point stack obtaining a time-lapse dataset of 270 frames. In order to follow the calcium dynamics over time in the elongating root hairs tips, registration of images was needed. The registration process was performed using the “napari-roi-registration plugin” (https://github.com/GiorgiaTortora/napari-roi-registration), developed for the multidimensional image viewer napari and based on the openCV python library, which allows to identify and follow an unlimited number of Regions of Interest (ROIs) simultaneously. In one frame of the 270-images stack, ROIs have been manually selected in correspondence of the root hair tips of interest and have been then registered automatically by the plugin. After registration, information related to the intensity and displacement of ROIs were extracted and saved using another feature of the napari-roi-registration plugin. Calcium variations over time were calculated from the raw intensities as the normalized R-GECO1 fluorescence intensity ΔF/F_0_ at each time point with the baseline F_0_ determined as the mean fluorescence value over time. Starting from the extracted normalized intensities, Fourier analysis was performed to determine the main oscillation frequencies in the detected calcium signals (Candeo et al., 2017). For each intensity signal, the fast Fourier transform was computed using a Python built-in function, in order to move from the time domain to the frequency domain and identify the characteristic oscillation frequency for each root hair. Fourier transform is a complex function composed of a real and an imaginary part, therefore, to identify the main oscillation frequencies, the power spectrum was extracted from the Fourier transform as:

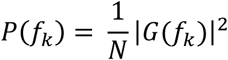

Where *N* is the number of time points and *G*(*f*_*k*_) is the Fourier transform defined in a discrete way as:

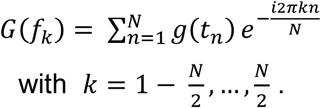

Finally, for each line group, the Power Spectral Density was computed as the mean of all the power spectra in that group and the error was calculated as the standard deviation. Kymograph images with median filter radius=1.0 were generated from MIP time-lapses (x, y, t) using the *Kymograph Builder* plugin by Fiji, to provide a space–time representation of selected pixels. Pixels were selected by tracing a line along the root hair, and a new image was generated, displaying the temporal variation in the intensities of the pixels beneath the selected line. Each column of pixels in the kymograph represents a single time point of the time-lapse dataset, while the vertical axis shows the intensities of the pixels along the line.

## Results

### *sune42* suppresses the dominant-negative effect mutant *LRX1ΔE14*

Expressing the truncated *LRX1* lacking the extensin domain (*LRX1ΔE14*) in wild-type Col induces aberrant root hairs that are misshaped, bulge, or burst (Figure 1A). *LRX1ΔE14* seedlings have very few root hairs that are elongated and their average length is shorter than Col (Figure 1B,C). This suggests that LRX1ΔE14 exerts a dominant-negative effect, likely through interfering with the process involving the endogenous LRX1. Hence, the *LRX1ΔE14* line is a genetic tool to investigate the physiological processes that are involved in the LRX1-related network. An ethyl methanesulfonate (EMS) mutagenesis was performed on the *LRX1ΔE14* line to screen for suppressors of the dominant-negative effect of *LRX1ΔE14* (Guérin et al., 2025). EMS-mutagenized *LRX1ΔE14* seeds were propagated to the M2 generation and M2 plants with reconstituted root hairs were selected for further analyses. Among those, *LRX1ΔE14 sune42* (***<u>su</u>****ppressor of dominant-**<u>ne</u>**gative effect 42*) displays a wild type-like root hair phenotype (Figure 1A). A more detailed quantification revealed that the root hair phenotype of *LRX1ΔE14* is largely but not completely suppressed by *sune42*, with root hairs that are slightly shorter and more misshaped compared to wild-type Col (Figure 1B,C). To exclude intragenic suppression of *LRX1ΔE14*, the construct *LRX1:LRX1ΔE14* was sequenced in *LRX1ΔE14 sune42* and revealed to be unaltered.

**Figure 1.**
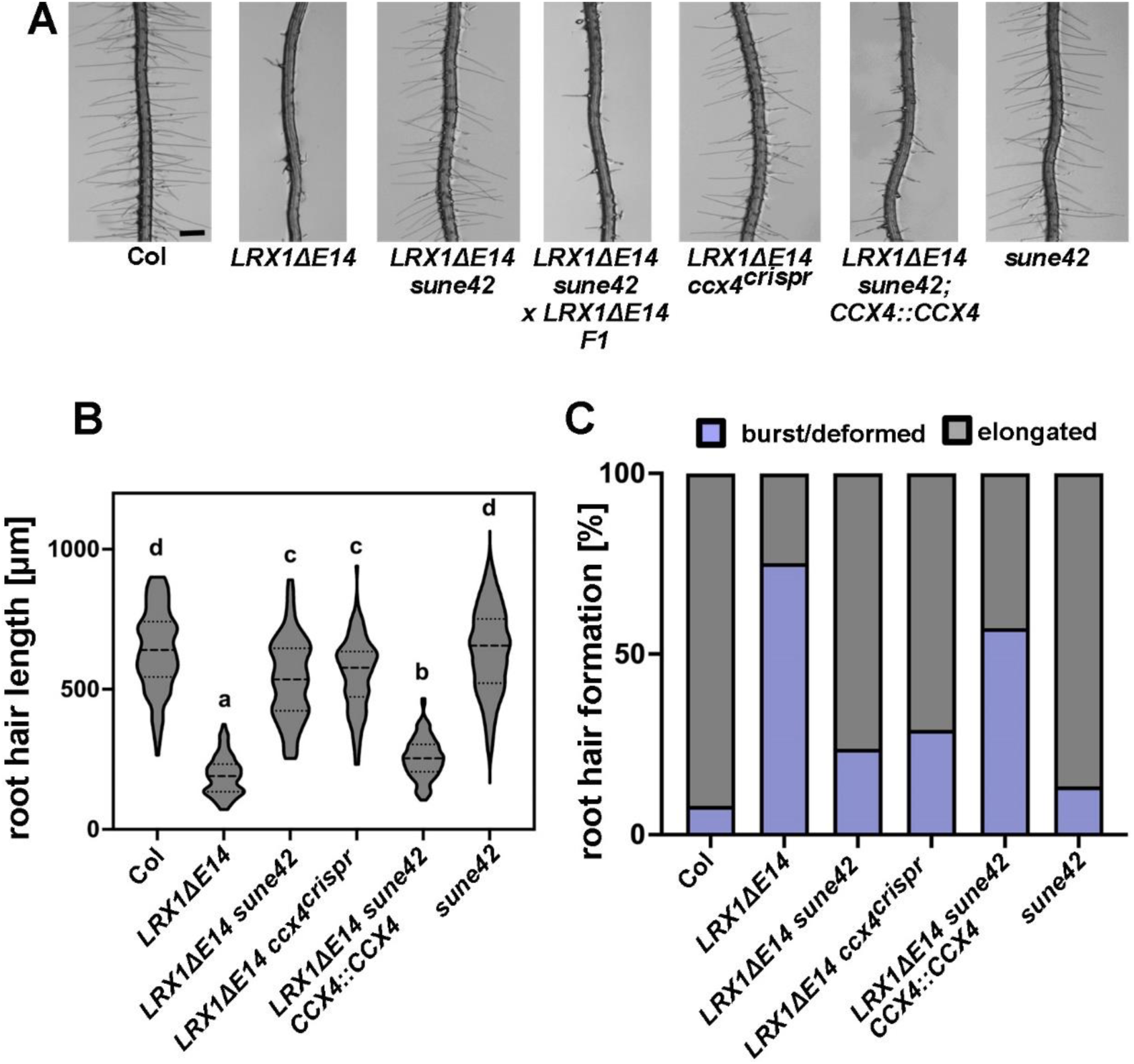
*sune42* suppresses the root hair defect of *LRX1ΔE14.* (A) Representative images of roots of 5-day-old *Arabidopsis* seedlings of the different lines. Seeds were germinated and vertically grown on ½ MS plates. Root hairs of *LRX1ΔE14* are severely defective compared to wild-type Col and restored by the *sune42* mutation, as well as the *ccx4^crispr^* mutation representing the *CRISPR/Cas9*-induced second allele of *ccx4.* The *sune42* single mutant shows a wild type-like root hair phenotype. *LRX1ΔE14 sune42 x LRX1ΔE14* is the F1 of a backcross confirming *sune42* being recessive and *LRX1ΔE14 sune42; pCCX4::CCX4* demonstrates complementation of *sune42* by the wild type copy of *CCX4*, inducing an *LRX1ΔE14*-like phenotype. Scale bar = 0.5 mm. (B) Violin plot shows the average length of elongated root hairs. Significant differences are indicated by letters (n>150 per genotype, one-way ANOVA with Tukey’s unequal N-HSD *post hoc* test, *P* < 0.01). Central dash line represents the median, dotted lines represent 25th and 75th percentiles (limits). (C) Stacked bar plot shows the percentages of root hairs with different shapes (n>150 per genotype).

A backcross of *LRX1ΔE14 sune42* with the *LRX1ΔE14* parental line revealed that *sune42* is a recessive mutation, as all the F1 seedlings exhibited a defective root hair phenotype resembling *LRX1ΔE14* (Figure 1A). Of a segregating F2 population, seedlings displaying a wild type-like *LRX1ΔE14 sune42* mutant phenotype were selected, pooled, and used for whole-genome sequencing (details see Materials and Methods). In comparison with *LRX1ΔE14*, which was previously sequenced (Guérin et al., 2025)*, LRX1ΔE14 sune42* revealed two non-silent single nucleotide polymorphisms (SNPs) in two nearby genes: in the gene coding for the putative cation/Ca Ca^2+^ exchanger *CCX4* (*At1g54115*) resulting in Gly187Glu (Suppl. Figure S1) and in the so far uncharacterized gene *At1g55750* (Ser502Leu). CAPS markers were established for the two mutations and tested on *sune42*-like seedlings of the segregating F2 population mentioned above. Among 82 F2 plants tested, only the SNP in *ccx4* showed complete linkage, suggesting that the *ccx4* mutation represents *sune42*. The *LRX1ΔE14 sune42* line was transformed with a wild-type *pCCX4::CCX4* construct, and several independent transgenic plants developed seedlings displaying *LRX1ΔE14*-like root hair development (Figure 1). Additionally, *CRISPR/Cas9*-mediated mutagenesis produced an additional *ccx4^crispr^* allele (Suppl. Figure S1) that also alleviates the *LRX1ΔE14* root hair defect (Figure 1), further confirming that *CCX4* represents the *sune42* locus.

A *sune42* single mutant was obtained by crossing *LRX1ΔE14 sune42* with wild-type Col and selecting for *sune42* and the absence of *LRX1ΔE14* in the segregating F2 generation. As shown in Figure 1, the *sune42* single mutant develops wild type-like root hairs, suggesting that *sune42* does not affect the root hair phenotype in the absence of *LRX1ΔE14.* The visual phenotype was confirmed to be equal to the wild type by quantifying root hair defects (Figure 1B,C).

### *sune42* partially suppresses the root hair defect in *lrx1 and fer-5*

The dominant-negative effect induced by *LRX1ΔE14* is likely due to its interference with the endogenous LRX1-RALF-FER network (Herger et al., 2020). Therefore, we examined the impact of the *sune42* mutation on LRX1 and FER functions in *lrx1 sune42* and *fer-5 sune42* double mutants. We observed partial alleviation of the typical *lrx1* and *fer-*5 root hair defects, with more elongated root hairs in *lrx1 sune42* and *fer-5 sune42* compared to *lrx1* and *fer-5*, respectively. Deformed root hairs, however, were still present in *lrx1 sune42* and *fer-5 sune42* (Figure 2). This suggests that *sune42* partially suppresses the *lrx1* and *fer-5* phenotypes.

**Figure 2.**
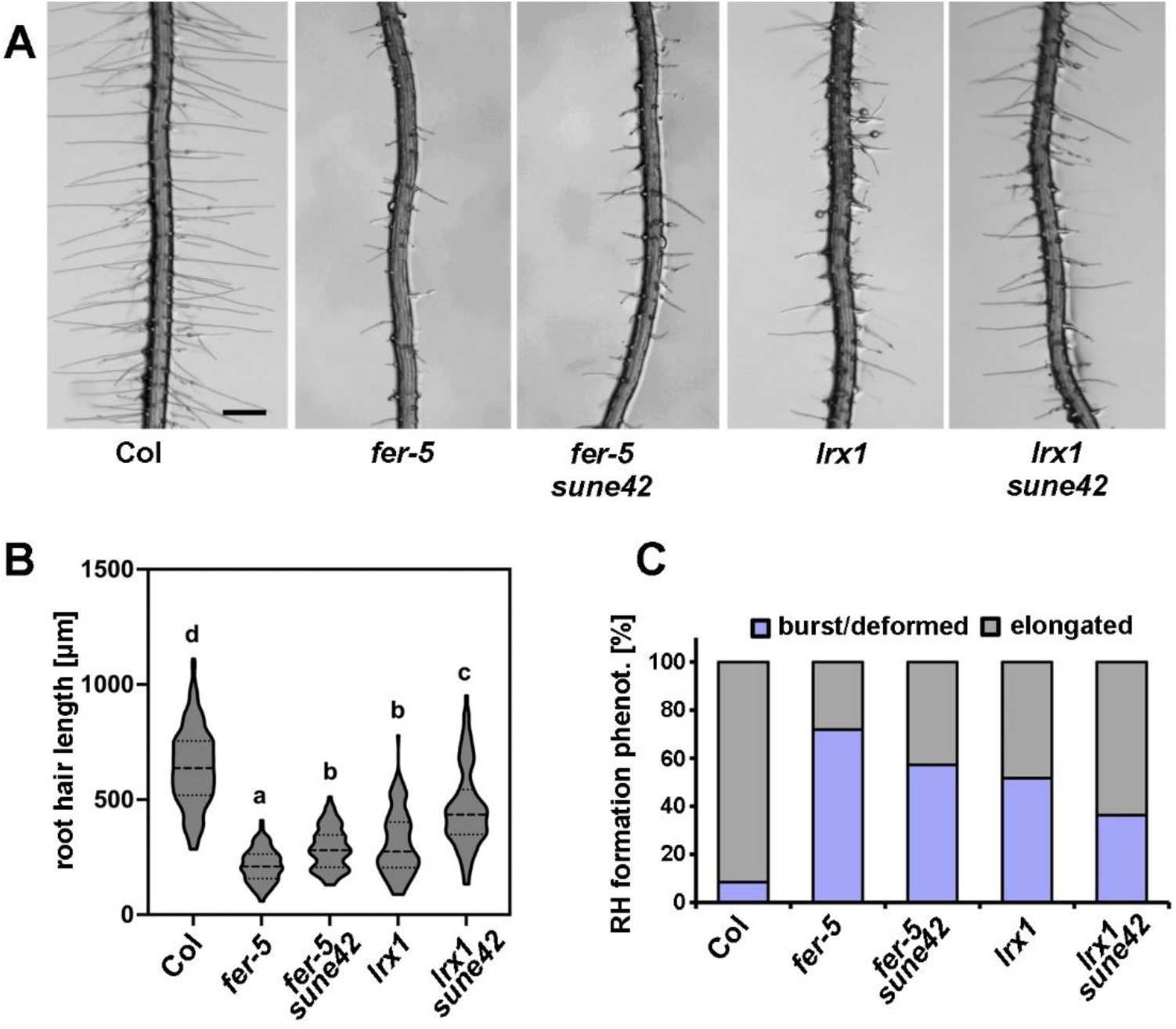
*sune42* partially suppresses *fer-5* and *lrx1* root hair phenotypes. (A) Representative images of 5-day-old *Arabidopsis* seedlings roots of Col, *fer-5*, *fer-5 sune42*, *lrx1*, and *lrx1 sune42.* All seedlings were grown vertically on ½ MS plates. *fer-5* and *lrx1* show aberrant root hair phenotypes. For both mutants, *sune42* results in slightly more/longer root hairs. Scale bar = 0.5 mm. (B) Quantification of the length of elongated root hairs is shown with the violin plot. Letters represent significant differences (n>150 per genotype, one-way ANOVA with Tukey’s unequal N-HSD *post hoc* test, *P* < 0.01). Central dash line represents the median, dotted lines represent 25th and 75th percentiles (limits). (C) The stacked bar plot shows the classification of root hairs with different shapes (n>150 per genotype).

### *sune42* alters salicylic acid signaling

It was previously reported that the loss of *CCX4* function leads to increased salicylic acid (SA) accumulation and expression of SA-responsive genes (Fujikura et al., 2020). Also, the LRX-FER module has been shown to modulate SA levels in the shoot (Zhao et al., 2021). With SA being important for root development and growth (Pasternak et al., 2019; Tan et al., 2020; Zhou et al., 2022), the involvement of SA signaling in the effect of *sune42* on root hair formation was investigated. First, SA levels were determined in Col, *LRX1ΔE14,* and *LRX1ΔE14 sune42* seedlings. Compared to *LRX1ΔE14*, the *sune42* mutation does not have a significant effect on SA levels in the root tissue but increases the SA levels in the shoot compared to *LRX1ΔE14*, in line with other findings (Fujikura et al., 2020) (Figure 3).

**Figure 3.**
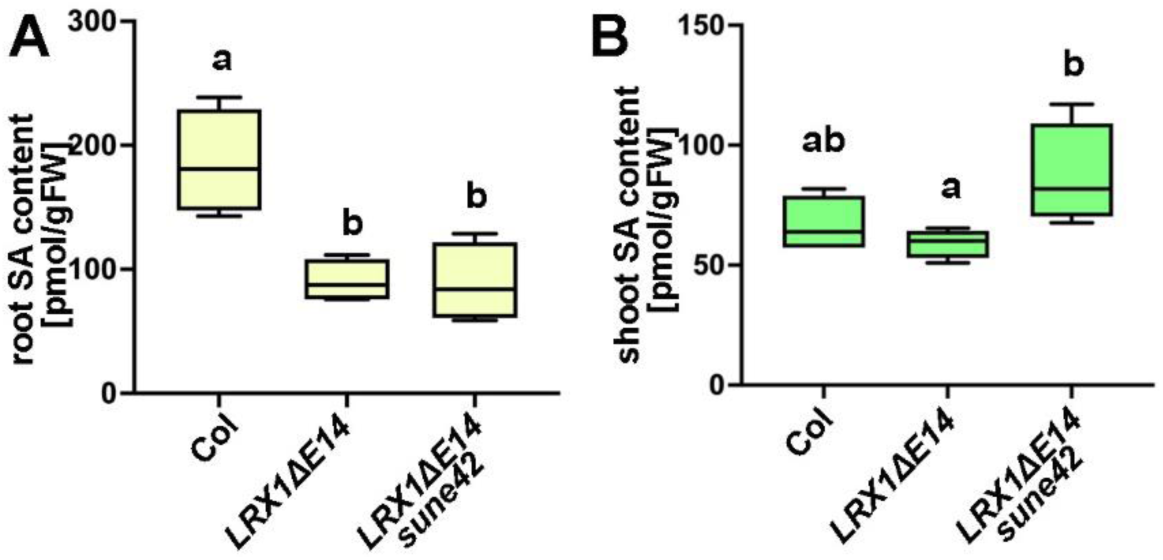
*sune42* causes increased shoot salicylic acid (SA) levels in *LRX1ΔE14*. Quantification of SA content in root (A) and shoot (B) tissues of 10-day-old *Arabidopsis* seedlings. The box plots represent the median (central line), 25th and 75th percentiles (limits), minimum/maximum values (whiskers), letters represent significance levels of statistical analyses (one-way ANOVA with Tukey’s unequal N-HSD *post hoc* test, *P* < 0.01, n=4). Plots showing primary data from one individual measurement.

Long-distance transport through the vascular network has been extensively studied for SA (Anfang and Shani, 2021), and it is possible that *sune42* causes accumulation of SA in the shoot that is transported to the root, where it promotes SA signaling and thereby influences root hair development. To explore if the root SA signaling is affected by the shoot SA levels, we measured SA-responsive gene expression in root tissue. qRT-PCR on mRNA extracted from seedling roots shows that the SA-responsive genes *WRKY70* and *WRKY53* (Yu et al., 2001; Li et al., 2004) are both downregulated in *LRX1ΔE14* in comparison with the wild type Col, and *sune42* rescues the gene expression level (Figure 4A), suggesting a possible correlation between the root hair phenotype and SA signaling.

**Figure 4.**
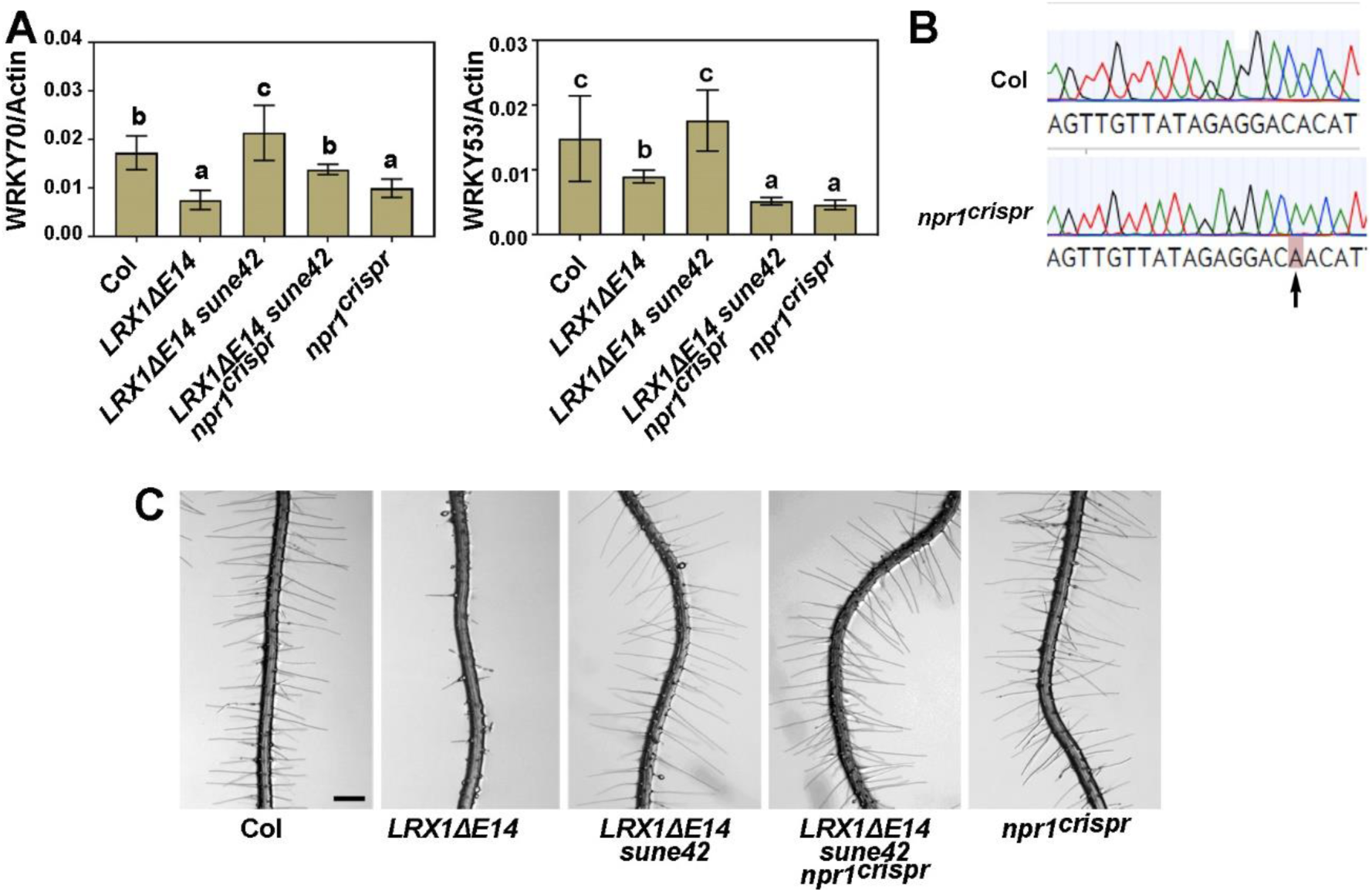
*sune42* suppresses *LRX1ΔE14* root hair phenotype independent of SA-induced NPR1 signaling. (A) qRT-PCR analyses showing transcript levels of SA responsive genes *WRKY53* and *WRKY71* in Col, *LRX1ΔE14, LRX1ΔE14 sune42, LRX1ΔE14 sune42 npr1,* and *npr1* lines. mRNA was extracted from 10-day-old *Arabidopsis* seedlings and reverse transcribed to cDNA for analyses. *ACTIN2* was used as an internal control. All values are normalized to *ACTIN2* transcript levels. Values are means of four biological replicates ± SD, letters indicate significance levels of statistical analyses (one-way ANOVA with Tukey’s unequal N-HSD *post hoc* test, *P* < 0.01) (B) Sequencing chromatogram showing the mutation of *CRISPR/Cas9*-mediated mutagenesis 603 bp downstream of the start codon. The insertion of one base pair at the codon Thr202 induces a frame shift and a stop codon after 8 amino acids. (C) Roots of 5-day-old vertically grown *Arabidopsis* seedlings. The root hair formation phenotype of *LRX1ΔE14* is suppressed by *LRX1ΔE14 sune42* and not further modified by *LRX1ΔE14 sune42 npr1^crispr^*. Scale bar=0.5 mm.

### *sune42* does not modulate *LRX1ΔE14* root hair defect through SA-dependent NPR1 signaling

To investigate the biological relevance of SA signaling in the *LRX1ΔE14-*induced root hair defect, we first tested the effect of exogenous SA on the root hair development in Col and *LRX1ΔE14*. As shown in Suppl. Figure S2, the presence of 10 μM SA did not cause any observable changes in Col and *LRX1ΔE14* in terms of root hair development compared to mock treatment. A mutation in *ccx4* has been shown to cause an SA-induced growth alteration that depends on the SA signaling component NPR1 (Wu et al., 2012; Fujikura et al., 2020). To test whether the *sune42*-induced effect on root hair formation is dependent on NPR1, a *CRISPR/Cas9*-mediated frame shift mutation in *NPR1* was induced in wild-type Col and *LRX1ΔE14 sune42* backgrounds, causing a single-nucleotide insertion 603 bp downstream of the start codon (Figure 4B). qRT-PCR shows that the expression of SA-responsive genes, *WRKY53* and *WRKY70,* is decreased in the *NPR1* loss-of function mutant background *npr1^crispr^*, in the context of the single mutant compared to Col as well as in the *LRX1ΔE14 sune42 npr1^crispr^* compared to *LRX1ΔE14 sune42* (Figure 4A). However, the root hair phenotype of *LRX1ΔE14 sune42* remains unaffected in the *LRX1ΔE14 sune42 npr1^crispr^* line (Figure 4C), implying that the *sune42*-mediated suppression of *LRX1ΔE14* is independent of the SA-dependent NPR1 signaling.

### Ca^2+^ deficiency promotes the root hair emergence in *LRX1ΔE14*

CCX proteins have been implicated in Ca^2+^ transport activities (Corso et al., 2018; Kanamori et al., 2023). On the other hand, mutations in *LRX* genes have been shown to be affected in Ca^2+^ homeostasis (Fabrice et al., 2018). This led us to hypothesize that the dominant-negative effect in *LRX1ΔE14* might be linked to impaired intracellular Ca^2+^ homeostasis, which would then be rescued by mutations in *CCX4*. To test that, we first investigated the effect of modified Ca^2+^ availability on the *LRX1ΔE14* root hair phenotype. Col and *LRX1ΔE14* seedlings were grown in the presence of the calcium chelator ethylene glycol-bis(β-aminoethyl ether)-N,N,N′,N′-tetra-acetic acid (EGTA). Root hair elongation was similarly inhibited in both Col and *LRX1ΔE14*; yet, *LRX1ΔE14* seedlings showed evidently more root hair emergence in the presence of 1 mM EGTA (Figure 5), which causes a reduction of free Ca^2+^ from 1.5 to 0.6 mM (https://somapp.ucdmc.ucdavis.edu/pharmacology/bers/maxchelator/CaMgATPEGTA-NIST.htm), indicating that limited Ca^2+^ availability rescues the root hair formation defect in *LRX1ΔE14* to a level that is close to the wild type.

**Figure 5.**
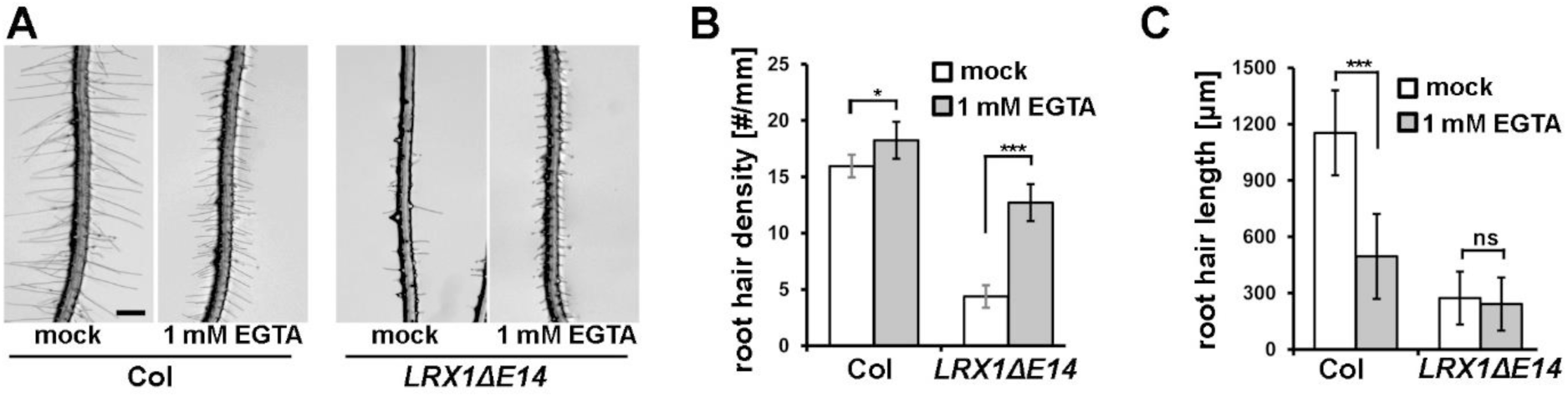
EGTA treatment leads to increased *LRX1ΔE14* root hair emergence. (A) Representative images of 5-day-old Col and *LRX1ΔE14* roots vertically grown on ½ MS plates with or without 1 mM EGTA. Scale bar = 500 μm. (B) Quantification of root hair density where only emerging root hair structures were counted (t-test, *:p<0.5, ***:p<0.0001, n>80 root hairs of four different seedlings). (C) Quantification of root hair length (t-test, ***:p<0.0001, ns: not significant, n>80 root hairs of four different seedlings).

### *sune42* stabilizes tip-focused Ca^2+^ oscillation in *LRX1ΔE14*

The EGTA assay implies that the dominant-negative phenotype of *LRX1ΔE14* is affected by the Ca^2+^ availability. To further understand how root hair development is affected by Ca^2+^, we examined the root hair tip-focused cytosolic Ca^2+^ dynamics by comparing Ca^2+^ oscillations in lines expressing the calcium indicator R-GECO1 driven by the *ubiquitin10* promoter (Keinath et al., 2015) by time-lapse single-cell Light Sheet Fluorescence Microscopy (LSFM) imaging. In this experimental set-up, the root hair phenotypes remained with the *LRX1ΔE14* showing impaired root hair formation, and alleviation of the root hair formation defect by *sune42* (Suppl. Figure S3). Kymographs were used to visualize temporal variations in the fluorescence signal. They provide a tool for a space-time representation from a time-lapse image sequence. In agreement with published data using the Cameleon YC3.6 Ca^2+^ indicator (Monshausen et al., 2008; Candeo et al., 2017), the tip-focused cytosolic Ca^2+^ of Col root hairs oscillates with a period which ranges from 18 up to 33 s (Figure 6A). Intriguingly, time-lapse experiments revealed altered Ca^2+^ oscillations in the root hairs of *LRX1ΔE14* (Figure 6A, Suppl. Video S1). While in root hairs of wild-type Col plants, a predominantly fast periodic oscillation is present, in *LRX1ΔE14* specimens a slow oscillation seems to be superimposed to the fast one. This effect is not evidently observable in *sune42* and *LRX1ΔE14 sune42.* To further compare the behaviour of the different populations, Fourier analysis was performed. The distribution of oscillation frequencies revealed a dominant frequency band extending from 0.057 to 0.03 Hz (from 18 to 33 s) in Col, while in *LRX1ΔE14* two distinct frequency bands were identified. In particular, a high frequency band similar to the main one of Col but slightly shifted towards lower values (from 0.050 to 0.025 Hz) is found together with a secondary drastically slower frequency band extending from 0.016 to 0.006 Hz (from 1 to 2.8 min) (Figure 6B), suggesting that *LRX1ΔE14* displays a combination of low and high Ca^2+^ oscillation frequencies (Figure 6B). By contrast, the line *LRX1ΔE14 sune42* shows a wild type-like Ca^2+^ oscillation pattern with a dominant frequency band extending from 0.055 to 0.025 Hz that overlaps with the frequency distribution of Col and a significant attenuation of the lower frequency band which also appears to be shifted to higher values (Figure 6B). Interestingly, the *sune42* single mutant shows only minimal alteration in Ca^2+^ oscillations compared to Col (Figure 6A,B). Taken together, our findings provide a correlation between root hair development and tip-focused Ca^2+^ oscillation that are both altered in *LRX1ΔE14* but reconstituted in *LRX1ΔE14 sune42*.

**Figure 6.**
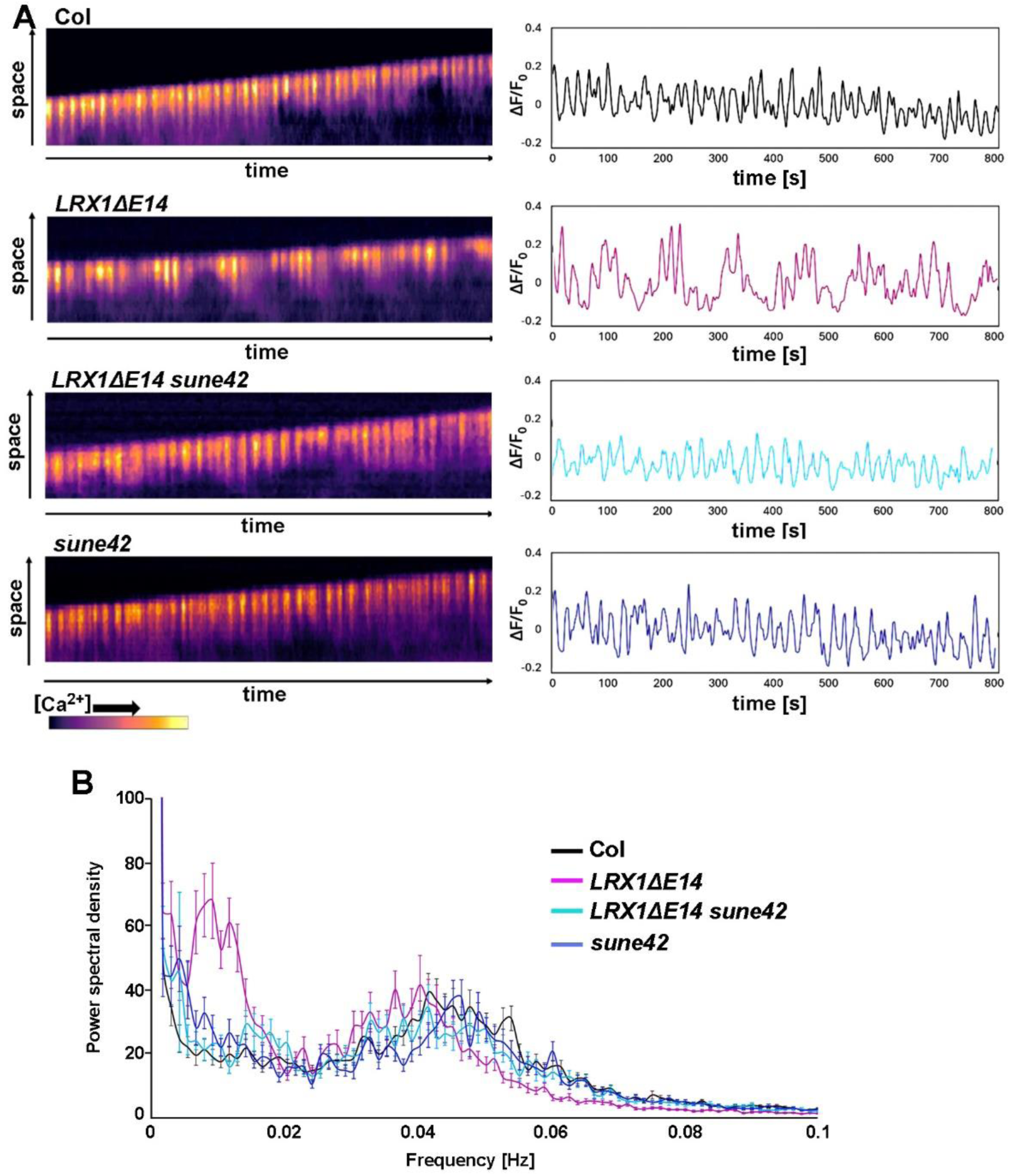
The *sune42* mutation alleviates the impaired tip-focused calcium oscillation in the root hairs of *LRX1ΔE14*. (A) Kymographs (left) and plotted data (right) of Ca^2+^ dynamics over time in growing root hairs of *R-GECO1*-expressing *Arabidopsis* seedlings of the different lines. Each line in the kymograph corresponds to a single time point in the time-lapse, while the vertical axis represents the intensity values. The root transition zone was used as normalized R-GECO1 fluorescence intensity (ΔF/F_0_). The sampling time was 3 seconds, and the measure was repeated 270 times. (B) Mean power spectral density obtained with Fourier transform spectral analysis. Dots of mean magnitude are connected to curves. Error bars are mean ± SD. (n=105 WT, 79 *LRX1ΔE14,* 67 *LRX1ΔE14 sune42,* and 75 *sune42* root hairs from more than 10 independent 5-day-old seedlings per genotype).

## Discussion

Mutations in *ccx4* cause a suppression of the dominant-negative effect of LRX1ΔE14 on root hair development. CCX4 was demonstrated to be involved in Ca^2+^ transport (Kanamori et al., 2023), and CCX2 revealed to be important for Ca^2+^ homeostasis (Corso et al., 2018). CCX4 contains two highly conserved Ca^2+^- and Na^+^-binding motifs, GNG(A/S)PD in α-1 and (G/S)(N/D)SxGD in α-2 (Suppl. Figure S1B), that were identified in the Na^+^/Ca^2+^ exchanger HsNCKX6 (Cai and Lytton, 2004), suggesting the potential role of CCX4 in Ca^2+^ transport. *sune42* possesses a missense mutation in the α-1 repeat of *CCX4*, resulting in Gly187Arg. Together, this points at Ca^2+^ fluxes being part of the LRX1ΔE14-mediated defect in root hair development, which is supported by the distorted cytosolic Ca^2+^ oscillations in root hairs of the LRX1ΔE14 line compared to the wild type. Ca^2+^ is a key modulator in controlling the polarity and the expansion rate of tip-growing cells (Monshausen et al., 2008; Candeo et al., 2017). Indeed, several cyclic nucleotide-gated channels (CNGCs) have been demonstrated to coordinate Ca^2+^ oscillations and root hair growth (Zhang et al., 2017; Brost et al., 2019; Zhu et al., 2025). Since *ccx4* alleviates both the modified cytosolic Ca^2+^ oscillations (Figure 6) and root hair growth defect in the *LRX1ΔE14* line (Figure 1), the correlation is established, but it does not prove that LRX1ΔE14 induces the altered Ca^2+^ dynamics. It is worth noting that mutations in pollen-expressed *LRX8,9,10,11* induce aberrant Ca^2+^ accumulation and limiting Ca^2+^ availability alleviates the growth defects in these mutants (Fabrice et al., 2018), suggesting a link between LRX-related processes and Ca^2+^ dynamics. The positive effect of the Ca^2+^ chelator EGTA on root hair formation (Figure 5) suggests that, also here, limiting Ca^2+^ availability helps overcoming the root hair defect by altered LRX activity. The activity of the Golgi-localized CCX4 tempers changes in Ca^2+^ dynamics (Kanamori et al., 2023) and appears to be involved in changing Ca^2+^ fluxes, comparable to the function of other members of the CCX family (Corso et al., 2018). This underlines the importance of organellar Ca^2+^ pools for regulating cytoplasmic Ca^2+^ homeostasis. In plants, a large amount of Ca^2+^ is bound in the cell wall via ionic interactions with galacturonic acid that is found in de-methylated pectin and Arabinogalactan proteins (Willats et al., 2001; Lopez-Hernandez et al., 2020). De-methylated pectin is bound by the LRX/RALF complex, which results in pectin compaction (Moussu et al., 2023; Schoenaers et al., 2024). It is conceivable that altering LRX/RALF/pectin interacting by modifying LRX availability has an impact on pectin dynamics and thus Ca^2+^ homeostasis, similar to the ability of RALF peptides to alter cytoplasmic Ca2+ concentrations (Haruta et al., 2008). A more detailed analysis of the impact of LRXs on pectin dynamics will possibly provide clues on the observation that *lrx* mutants (Fabrice et al., 2018) and LRX1ΔE14 impact Ca^2+^ dynamics.

The *xs2* mutant is an allele of *ccx4* and develops a defect in cell expansion due to the overaccumulation of SA signaling (Fujikura et al., 2020). This point needed to be further investigated, since (Zhao et al., 2021) showed that the salicylic acid (SA) and jasmonic acid (JA) pathways are constitutively upregulated in the *lrx3 lrx4 lrx5* mutant, providing a potential link between LRX activity and SA signaling. Our data show that SA-responsive genes are downregulated in *LRX1ΔE14* but upregulated again by the *sune42* mutation, as is the SA level in the shoot (Figure 3 and Figure 4), confirming the influence of SA signaling by *ccx4*. While a mutation in *NPR1* (Wu et al., 2012), the key responsive regulator of SA signaling, rescues the cell expansion defect in *xs2* shoots (Fujikura et al., 2020), our results show that a mutation in *NPR1* does not influence the root hair phenotype of *LRX1ΔE14 sune42* (Figure 4). This suggests that the effect of *sune42* on the *LRX1ΔE14*-mediated root hair defect is independent of the altered SA signaling.

The effect of the *ccx4* mutations is likely to be multifaceted. The Golgi apparatus is not only an organelle with high Ca^2+^ concentration that helps regulating cytoplasmic [Ca^2+^] (Ordenes et al., 2012), but it is also central to posttranslational protein modifications and the synthesis of numerous polysaccharides destined for the cell wall (Cosgrove, 2024). Glycosylation is the major intracellular posttranslational modification of cell wall proteins (Nguema-Ona et al., 2014). In mammalian cells, these processes are influenced by Ca^2+^, which modifies vesicle transport dynamics (Gerdes et al., 1989) and many glycosyltransferases and glycosidases involved in glycosylation (Bai et al., 2006). Altering Golgi-localized processes can thus lead to changes in the cell wall. *rol16* (*repressor of lrx1_16*) is affected in the Golgi-localized Apyrase 7 (Gupta et al., 2024), confirming that the effect of modified LRX1 function can be influenced by processes emanating from the Golgi.

LRX proteins, in conjunction with the plasma membrane-localized receptor kinase FER, influence vacuole development and cell growth, resulting in comparable mutant phenotypes of *fer* and higher-order *lrx* mutants (Dünser et al., 2019; Herger et al., 2020). While *sune42* has a strongly alleviating effect on *LRX1ΔE14* (Figure 1), its impact on the *lrx1* and *fer-5* mutant phenotypes is limited (Figure 2). This suggests that the *LRX1ΔE14* dominant-negative and the *lrx1* loss-of-function mutant have different causes. Interestingly, the truncated, extensin-less, variant of the root/shoot-expressed *LRX4* (Draeger et al., 2015), *LRX4ΔE*, also induces a dominant-negative phenotype, but at the same time renders the plant hypersensitive to RALF peptides (Dünser et al., 2019). These plant peptide hormones are bound by LRX proteins (Mecchia et al., 2017; Moussu et al., 2020), impact pectin compaction, and inhibit root growth (Abarca et al., 2021; Moussu et al., 2023; Schoenaers et al., 2024). This suggests that extensin-less LRXs, *LRX4ΔE* and *LRX1ΔE14*, are present but not anchored in the cell wall, changing their spatial distribution and making them available for interaction partners at incorrect locations. Thus, while the *LRX1ΔE14*-induced root hair phenotype is comparable to a mutant lacking root hair-expressed *LRXs* (Baumberger et al., 2003), LRX1ΔE14 actively interferes with the LRX1-related process in root hairs. The first identified *sune* mutant, *sune82*, is affected in His biosynthesis and impacts the TOR (Target of Rapamycin) network (Shi et al., 2018; Guérin et al., 2025). In contrast to *sune42* characterized here, *sune82* also suppresses *lrx1* (Guérin et al., 2025). Thus, it is important to use several approaches to identify suppressor mutants with different targets in the processes modified by LRX1. Depending on where in the process they take influence, these suppress only one or both types of distorted LRX1 activity. This will help us better understand the plethora of cellular events involving LRXs.

## Acknowledgements

We thank the infrastructure team at IPMB was continuous support with growth facilities. This work was supported by the University of Zurich and the Swiss National Science Foundation grants Nr 31003A_166577/1 and 310030_192495 to CR; the Ministero dell’Istruzione, dell’Università e della Ricerca — Fondo per Progetti di Ricerca di Rilevante Interesse Nazionale 2022 to A.C. and S.B; the Agritech National Research Center, funded by the European Union NextGenerationEU (Piano Nazionale di Ripresa e Resilienza (PNRR) — Missione 4, Componente 2, Investimento 1.4 – D.D. 1032 17/06/2022, CN00000022) to A.C.; and the Fondazione Fratelli Confalonieri to S.B..

## Author contributions

XH: *CRISPR/Cas9* mutagenesis, *CCX4* complementation tests, salicylic acid assays, Ca^2+^ deficiency assays

GT: Ca^2+^ oscillation measurements

AH: EMS mutagenesis, *sune* mutant screen

SB: Ca^2+^ oscillation measurements

PD: Hormonal content analysis

RV: Hormonal content analysis

JL: Hormonal content analysis

AS: whole-genome sequencing

GK: genetic analysis

MS: *sune* mutant analysis

AC: Ca^2+^ measurements

TW: whole-genome sequencing, supervising

AC: Ca^2+^ measurements, supervising

AB: Ca^2+^ measurements, supervising

CR: conceptualizing research, supervising, funding

All authors contributed to the writing process

## Data availability

All data are shown. Raw data will be made available upon request.

## Supporting information

**Supplemental Figures 1-3** are part of this manuscript and shown below.

**Supplemental Video S1** Video of Ca^2+^ oscillations, monitored by R-GECO1 fluorescence in the four genotypes wild type (Col), *LRX1ΔE14*, *LRX1ΔE14 sune42*, and *sune42*. These are accessible as a separate file accompanying this manuscript.

**Supplemental Figure S1.**
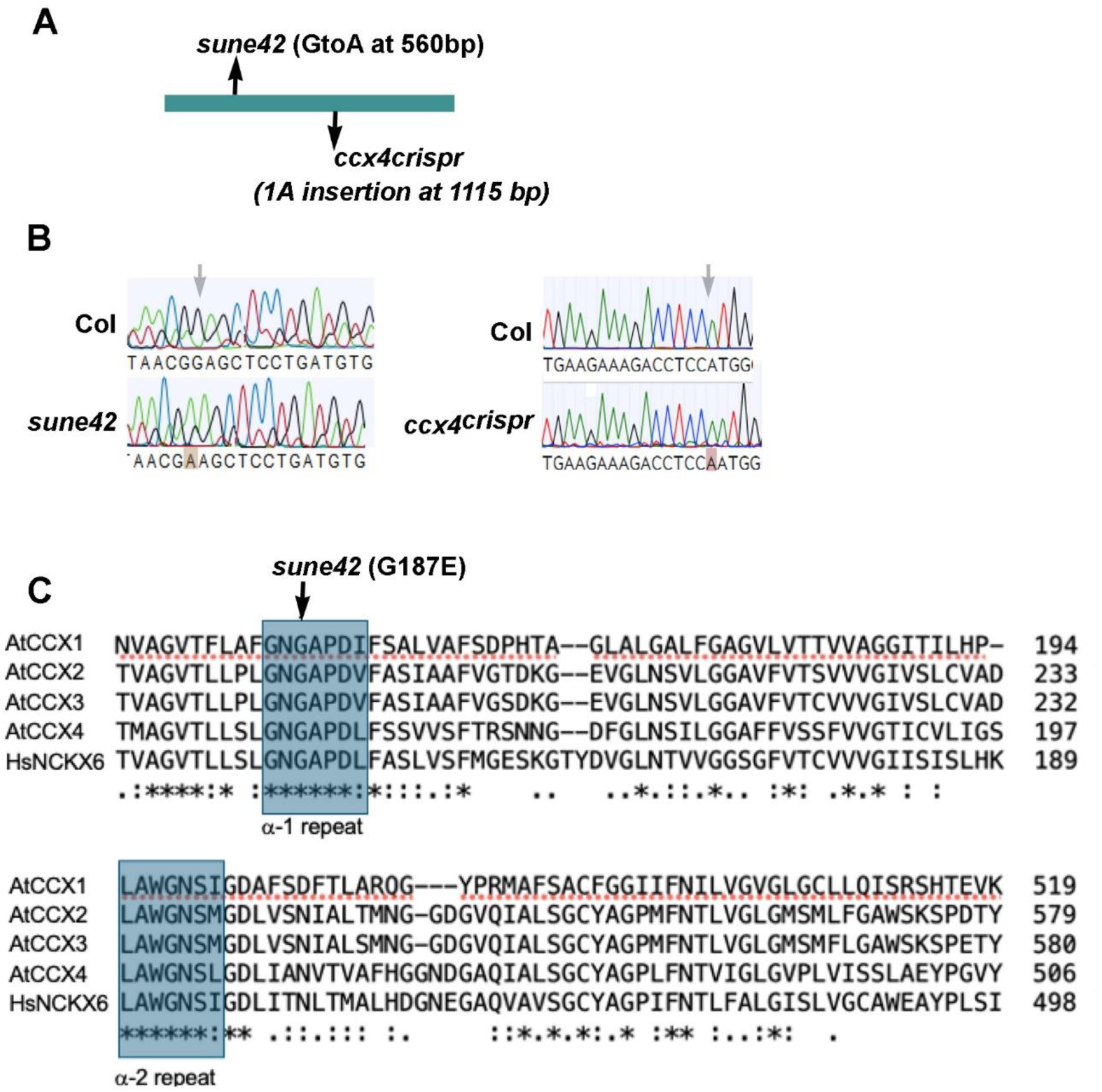
Scheme of *CCX4* genomic sequences and amino acid sequence alignment. (A) Schematic diagram of the intronless *CCX4* (*At1g54115*) genomic sequence of 1935 bp, showing the missense mutation site of the EMS allele *sune42* (G to A mutation at position 560 from the start codon resulting in a Gly187Glu substitution) and the *CRISPR/Cas9*-guided insertion of one A at position 1115 relative to the start codon that leads to a shift in the ORF after Pro372 and an early stop codon after 48 base pairs. (B) Sequencing chromatograms of the wild-type, *sune42*, and the *CRISPR/Cas9*-induced *ccx4* allele. Arrows indicate altered positions, brownish colorings show altered sequences. (C) Amino acid sequence alignment of CCX4 with its homologs from *Arabidopsis* AtCCX1, AtCCX2, AtCCX3 and its ortholog from *Human* HsNCKX6. Blue boxes indicate the conserved α repeats. The alignment was done with Clustal Omega (https://www.ebi.ac.uk/jdispatcher/msa/clustalo) (Sievers et al., 2011). Identical, conserved, and similar positions in the alignment are indicated by asterisks, colons, or single dots, respectively.

**Supplemental Figure S2.**
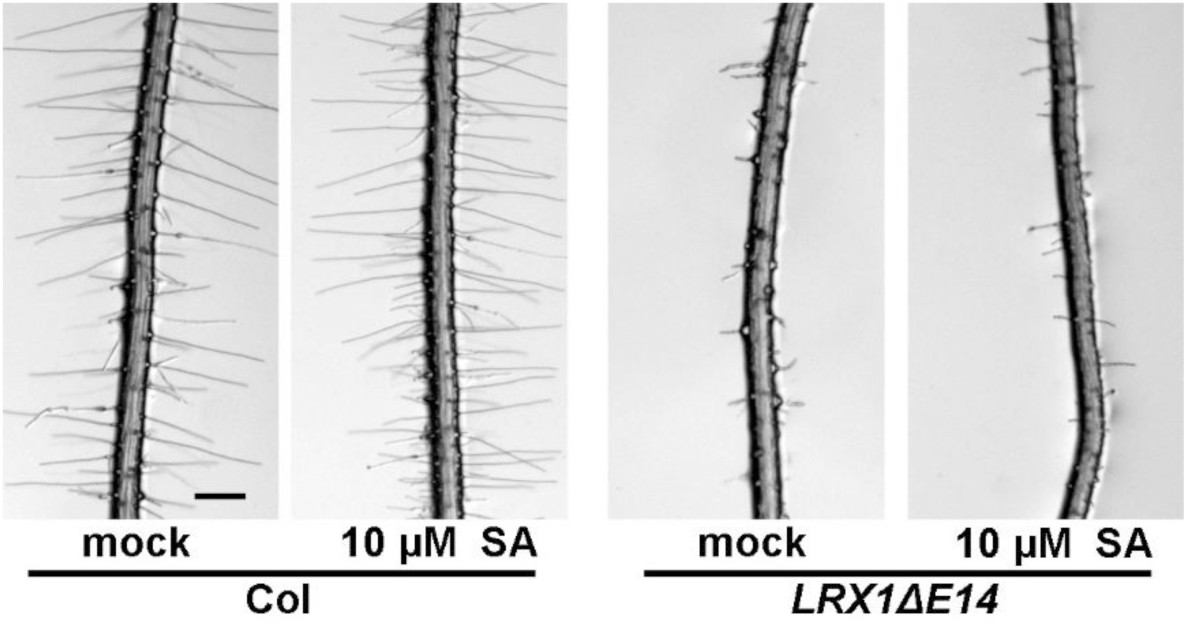
Exogenous SA does not affect the root hair development. Col and *LRX1ΔE14 Arabidopsis* seedlings were grown on ½ MS plates with or without 10 μM SA for 5 days in a vertical orientation. Representative images of mock (left) and 10 μM SA (right) of each genotype are shown in pairs. Both Col and *LRX1ΔE14* do not show altered root hair phenotype upon 10 μM SA treatment. Similar results were observed among three independent experiments.

**Supplemental Figure S3.**
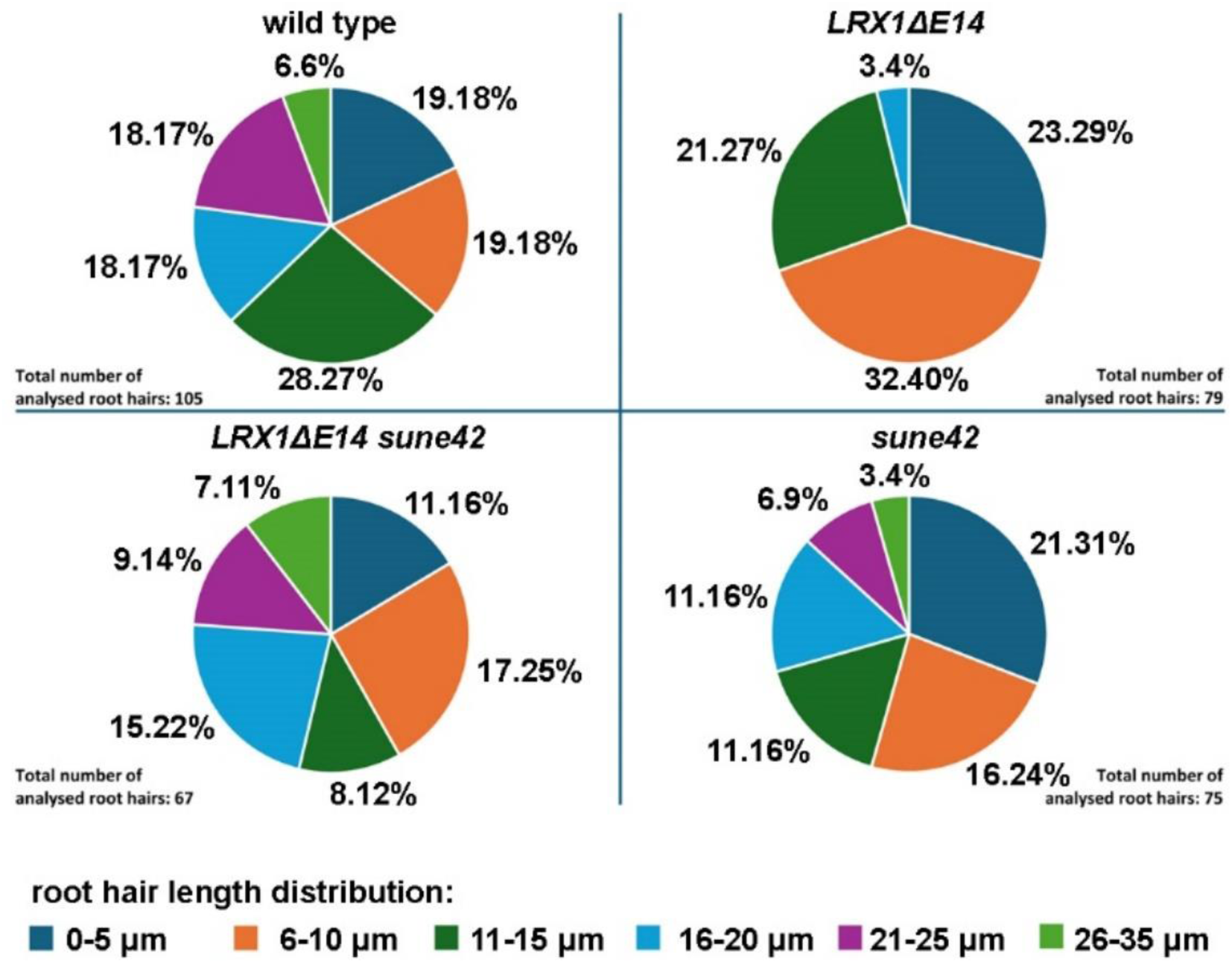
Difference in root hair formation between genotypes are maintained under LSF microscopy. For Light Sheet Fluorescence (LSF) microscopy analysis, seedlings needed to be growth into the agar. Under these conditions, the *LRX1ΔE14* line still showed strongly impaired root hair formation compared to the wild type, and this effect is suppressed by *sune42*. The pie-charts depict root hair length distributions in the different lines (n>66). Root hairs longer than 21 μm (purple and pale green sectors) in the wild type or in the presence of *sune42* were not detected in *LRX1ΔE14*.

